# A novel approach to evaluate alpha-synuclein seeding shows a wide heterogeneity in multiple system atrophy

**DOI:** 10.1101/2021.08.10.455800

**Authors:** Ivan Martinez-Valbuena, Naomi P. Visanji, Ain Kim, Heather H. C. Lau, Raphaella W. L. So, Sohaila Alshimemeri, Andrew Gao, Michael Seidman, Maria R. Luquin, Joel C. Watts, Anthony E. Lang, Gabor G. Kovacs

## Abstract

Several *in vitro* and *in vivo* findings have consistently shown that α-synuclein derived from multiple system atrophy (MSA) subjects has more seeding capacity than Parkinson’s disease-derived α-synuclein. However, reliable detection of α-synuclein derived from MSA using seeded amplification assays, such as the Real-Time Quaking-induced Conversion, has remained challenging. Here we demonstrate that the interaction of the Thioflavin T dye with α-synuclein from MSA and Parkinson’s disease patients can be modulated by the type of salt, pH, and ionic strength used to generate strain-specific reaction buffers. Employing this novel approach, we have generated a streamlined Real-Time Quaking-induced Conversion assay capable of categorizing MSA brains according to their α-synuclein seeding behavior, and to unravel a previously unrecognized heterogeneity in seeding activity between different brain regions of a given individual that goes beyond immunohistochemical observations and provide a framework for future molecular subtyping of MSA.

## Introduction

Multiple system atrophy (MSA) is a progressive neurodegenerative disorder with a clinical presentation of various combinations of parkinsonism, cerebellar and autonomic dysfunction^1^. Patients with predominant parkinsonian features are designated MSA-P, and patients with predominant cerebellar ataxia as MSA-C; however, the predominant motor feature can change over time^1^. The term MSA was coined in 1969 to pool previously described neurological entities^2^, however, the major common finding of argyrophilic oligodendrocytic cytoplasmic inclusions (GCIs), called Papp-Lantos bodies contributed significantly to the nosological definition of the disease^3^. Presence of the 140 aa protein, α-synuclein as a major component of these inclusions linked MSA with Lewy body disorders such as Parkinson’s disease (PD) and dementia with Lewy Bodies (DLB)^4^. Collectively, these disorders are now termed synucleinopathies. Lewy body disorders are classically distinguished from MSA by distinct cellular pathology. Although both conditions accumulate α-synuclein in a variety of cell types, MSA is characterized by oligodendrocytic inclusions, while neuronal α-synuclein pathology predominates in Lewy body disorders. In addition, recent studies have uncovered the presence of α-synuclein “polymorphs”, suggestive of different strains as occurs in prion diseases, that might be the basis for the phenotypic diversity found in these conditions. The presence of different strains is hypothesized to dictate the cell-to-cell spreading of pathology and the cellular impact of the pathological α-synuclein in every individual^5,6^. Consistent with this notion, many experimental findings have indicated that α-synuclein forming GCIs has greater seeding activity compared to Lewy body associated α-synuclein^7^. In seeding studies using cultured cells, GCI-α-synuclein was found to be approximately 1,000-fold more potent than LB-α-synuclein in triggering *de novo* α-synuclein aggregation^8^. Furthermore, recent discoveries using cryo-electron microscopy showed structural differences between the aggregates found in MSA and DLB brains^9,10^. In spite of these observations, on a neuropathological level, MSA is still defined by the presence of α-synuclein positive GCIs associated with neurodegenerative changes in striatonigral or olivopontocerebellar structures^11^. Moreover, although the distribution of inclusions correlates with the predominant clinical features^12,13^, there are currently no cytopathological or biochemical features for further subtyping of diverse MSA cases.

In recent years, the use of seeded amplification assays has emerged as a reliable method of detecting minute amounts of misfolded disease-associated proteins or seeds such as prion protein, tau or α-synuclein^14^. These assays, also known as Real-Time Quaking-induced Conversion (RT-QuIC) and protein misfolding cyclic amplification (PMCA), exploit the property of self-propagation to amplify and then sensitively detect minute amounts of these protein seeds^15,16^. Real-time detection of thioflavin T (ThT) fluorescence at multiple timepoints during the assay permits the measurement of kinetic differences between the seeding properties of different samples, providing a highly specific characterization of whether a disease-associated protein is present or not. However, despite the reliability of these assays to consistently detect α-synuclein seeds in Lewy body disorders, using a variety of biological samples^17–20^, attempts to detect α-synuclein in MSA have proven challenging. Van Rumund *et al*. found that α-synuclein RT-QuIC was positive in only 6/17 CSF samples in MSA^19^, while Rossi *et al*. detected an even lower number of RT-QuIC positive CSF samples in MSA (2/29)^20^. In contrast, using PMCA over a period of 350 hours, Shahawanaz *et al*. were able to detect α-synuclein seeding activity in 65/75 MSA CSF samples, but noted that in spite of aggregating faster, MSA CSF and brain samples reached a lower fluorescence plateau than PD CSF and brain samples^21^. These divergent findings are likely the result of different conditions in the reaction buffers used in these studies. Thus, in the present study, we sought to identify the optimal assay conditions that favor MSA seeding activity, by systematically modulating the ionic and cationic composition and the pH of the α-synuclein RT-QuIC reaction buffer. The effect of ionic composition and pH in α-synuclein aggregation has been widely studied^22^, but not in the context of the systematic comparison of a spectrum of reaction buffers seeded with brain homogenates from MSA and PD. Our analysis of 168 different buffer conditions revealed that changes in the physicochemical factors that govern the *in vitro* aggregation of α-synuclein induced strikingly divergent aggregation patterns between the PD and MSA samples examined. Further, employing this novel approach, we demonstrate a previously unrecognized heterogeneity in the 1) inter-individual seeding behavior of α-synuclein obtained from different MSA patients and, 2) intra-individual heterogeneity, with seeding activity varying between different brain regions of a given individual. Our findings go beyond any differences previously described using conventional immunohistochemistry and provide support for the future molecular subtyping of MSA.

## Results

### Screening 168 RT-QuIC conditions to detect α-synuclein seeding in MSA

We hypothesized that changing the physicochemical factors that govern the *in vitro* amplification of amyloidogenic proteins would favor α-synuclein seeding in MSA. Based on the published success of using different ionic environments to enhance the sensitivity of different proteopathic seeding amplification assays^22^, we conducted a systematic evaluation of 168 different reaction buffers, using an array of pH and salts, seeded with brain homogenates from one MSA and one PD patient (Fig. 1). We compared the effects of conducting the α-synuclein RT-QuIC in the presence of strongly (Citrate^3-^ and S_2_O_3_^2-^), moderately (F^-^ and Cl^-^) and weakly (ClO_4_^-^) hydrated anions and cations (guanidine hydrochloride (GdnCl) and MgCl_2_) at high (500 and 350 mM), medium (250 and 170 mM) and low (100 and 50 mM) concentrations using 4 different pH (pH2, pH4, pH6.5 and pH8). To seed the resulting 168 buffers, equal amounts (5μg) of total brain protein were used per reaction. A region rich in Lewy body pathology, the substantia nigra (SN), from a PD patient and a region enriched with GCIs, the cerebellar white matter, from an MSA patient were carefully microdissected using a 3 mm biopsy punch and subsequently processed into crude homogenates, using PBS. In these two cases, conventional immunohistochemistry for disease-associated α-synuclein demonstrated abundant Lewy bodies and neurites in the SN of the PD patient and GCIs in the cerebellar white matter of the MSA patient (Extended data Fig. 1a-b). Next, we looked for further biochemical and structural differences in the α-synuclein brain homogenates and using Western blotting found that the PD extract exhibited a different detergent-insoluble phosphorylated α-synuclein banding pattern, with more high-molecular-weight species than the MSA extract (Extended data Fig. 1c). In addition, the banding pattern of thermolysin-resistant α-synuclein species in MSA was different from that in PD. While both brain extracts possessed a band at ∼10 kDa, the MSA extract displayed a prominent additional band at ∼15 kDa (Extended data Fig. 1d). Finally, using a conformational stability assay (CSA), that measures the differential abilities of protein aggregate strains to resist denaturation by GdnCl, we found that α-synuclein aggregates from the MSA brain extract were significantly less stable than those present in the PD brain extract (Extended data Fig. 1e, f). Taken together, these results demonstrate that the MSA and the PD brain extracts used to seed our reactions contained conformationally distinct species of α-synuclein.

**Figure 1.**
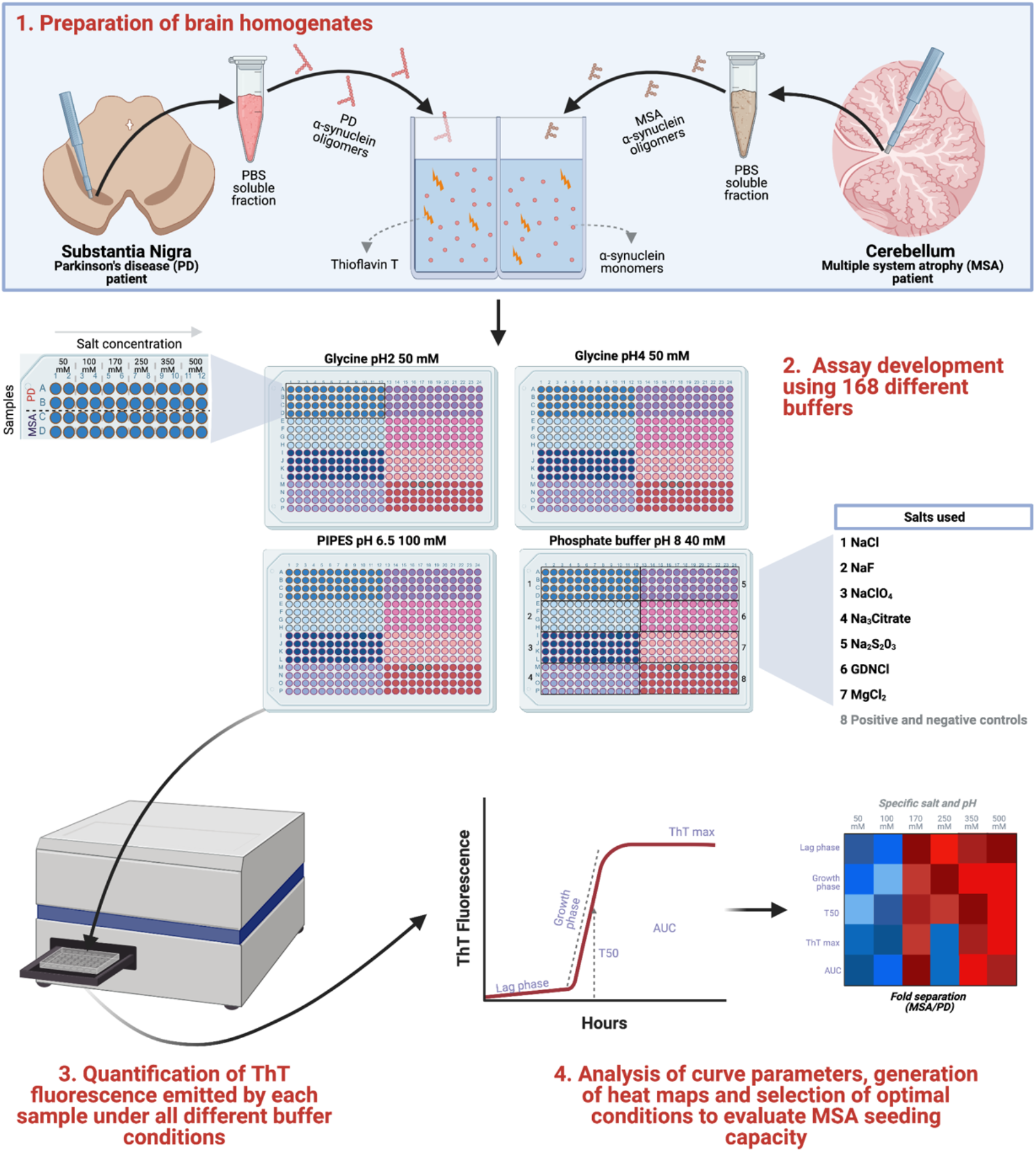
Schematic of the RT-QuIC buffer discovery phase to evaluate α-synuclein seeding capacity in MSA. 1.1) Samples containing α-synuclein seeds were prepared from substantia nigra of PD or cerebellum of MSA patients. 1.2) Samples were incubated with 168 reaction buffers and subjected to cycles of shaking and resting at 37ºC. 1.3) ThT output was measured at multiple timepoints over 48 hours. 1.4) Results were analyzed and heatmaps were generated to determine the optimal conditions to discriminate MSA-derived samples from PD-derived samples.

Following the characterization of the brain homogenates that would be used to seed the reactions, the 168 different conditions were run in quadruplicate and the same positive and negative controls were added in each plate. We quantified 5 kinetic parameters from each reaction: lag time, growth phase, T50 (which corresponds to the time needed to reach 50% of maximum aggregation), ThT max or fluorescence peak, and area under the curve (AUC) of the fluorescence response. Importantly, we confirmed that no significant differences were observed in any of these 5 parameters between the positive and negative control samples run on all plates. Next, we calculated the fold separation values of the 5 parameters between the curves obtained from the MSA and the PD patients, and generated heatmaps (Fig. 2). To select the conditions that were most favorable to detect MSA-derived α-synuclein aggregation in comparison with PD-derived α-synuclein we defined the AUC as the seeding parameter of interest as it incorporates all the kinetic features of each aggregation reaction, including the speed and extent of aggregation. Our analysis revealed that 19/168 buffers were able to detect predominantly PD-derived α-synuclein aggregation, showing at least twofold difference between PD and MSA curves. 109/168 buffers showed no major differences in the main parameters describing the kinetic curve of the RT-QuIC when the MSA aggregation pattern was compared to PD. 40/168 buffers showed at least twofold difference between MSA-derived α-synuclein and PD-derived α-synuclein aggregation. From these 40 buffers, we carefully examined the parameters describing the kinetic curves from the 10 conditions that displayed the highest differences. Based on the raw values obtained with each condition, we selected two buffers for further analysis.

**Figure 2.**
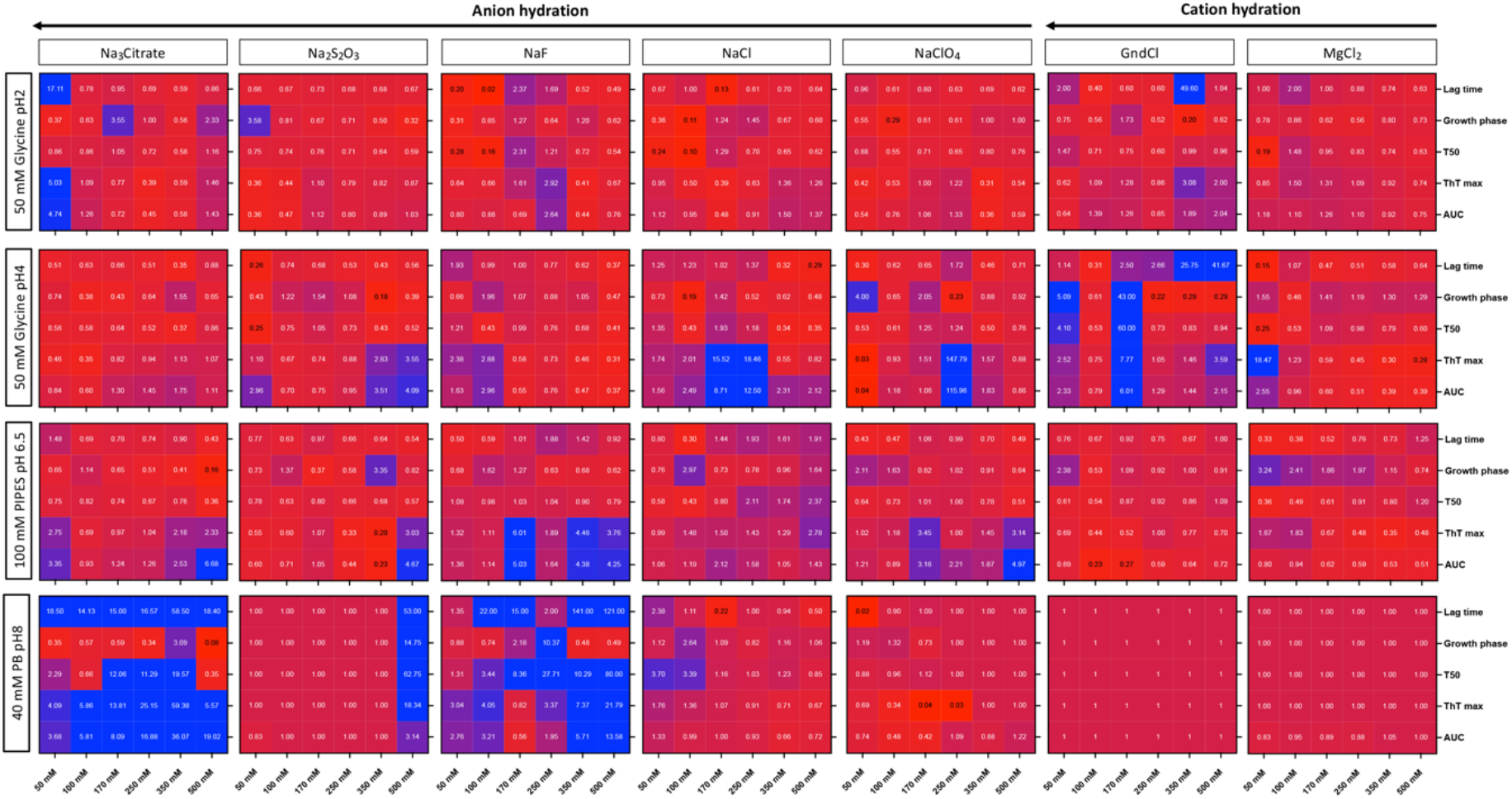
Changes in RT-QuIC physicochemical factors promote the detection of α-synuclein seeding capacity in MSA. Fold separation of lag time, growth phase, T50, ThT max and area under the curve (AUC) between reactions seeded with MSA-or PD-derived α-synuclein aggregates and amplified under 4 different pH using 7 salts at 6 concentrations. Larger numbers (blue) represent the conditions that showed a more favorable environment to detect MSA α-synuclein aggregation. When the fold separation is 1 (deep red), no discrimination between MSA and PD aggregation was found. Lower numbers (<1, lighter red/purple) represent the conditions more favorable to detect PD aggregation over MSA. *Buffers highlighted in yellow boxes represent the optimal conditions to detect α-synuclein seeding capacity in MSA and discriminate from PD.

### Validation of the two optimal conditions for assessing α-synuclein seeding in MSA

Having selected the two optimal buffers to detect MSA-derived α-synuclein seeding activity and differentiate it from PD-derived α-synuclein seeding activity, we proceeded to validate the results in a larger cohort of disease samples. We microdissected the SN of 15 subjects with MSA, 15 subjects with PD, 5 subjects with supranuclear progressive palsy (PSP) and 5 controls and processed the samples into PBS soluble homogenates. All subjects had a detailed neuropathological examination (Extended data Table 1). Equal amounts (5μg) of total protein were used to seed the reactions from the 42 subjects, using the two buffer conditions, with all samples run in quadruplicate on the same plate. To illustrate the typical profile of α-synuclein RT-QuIC aggregation for samples of PD and MSA, we plotted data from one representative MSA, one PD and one control case (the median subject was used in each example curve) amplified under the buffer 1 condition (Fig. 3a). The maximum fluorescence and the AUC were consistently different for PD and MSA, with samples from patients with MSA consistently aggregating faster while always reaching a lower fluorescence plateau than samples from patients with PD (Fig. 3 b-c). When we plotted data from the same cases but amplified under the buffer 2 condition, we observed that samples from patients with MSA still aggregated faster but in contrast, using this buffer, they reached a higher fluorescence plateau than those from patients with PD (Fig. 3e). Furthermore, significant differences in the maximum fluorescence and the AUC were observed between PD and MSA patients (Fig. 3f-g). To test whether the amount of ThT bound to the MSA and PD fibrils depends on the physicochemical factors of the buffer, we investigated if the end products from the RT-QuIC assay amplified under the two conditions showed structural differences when examined using electron microscopy (Fig 3 d, h). Interestingly, and despite the striking differences in the ThT kinetics shown by the two buffers, under both conditions the fibrils derived from the PD samples consistently showed a thin and straight morphology, while the fibrils derived from the MSA samples were consistently thicker and twisted. When we compared the kinetics shown by the PD and MSA patient samples with a cohort of controls and PSP patient samples without α-synuclein co-pathology, the MSA samples did not show significant differences in either the maximum fluorescence or in the AUC compared to controls or PSP subjects amplified under the buffer 1 condition (Fig. 3b-c). However, the MSA samples showed faster aggregation (T50) than the control and PSP group (Fig. 3a). When the buffer 2 condition was examined, we found an optimal distinction between MSA and PD, and also maximized the kinetic separation from controls and PSP subjects both in the maximum fluorescence and in the AUC (Fig. 3f-g). Therefore, the buffer 2 was selected as the optimal working buffer to study MSA α-synuclein seeding in RT-QuIC assays.

**Fig 3.**
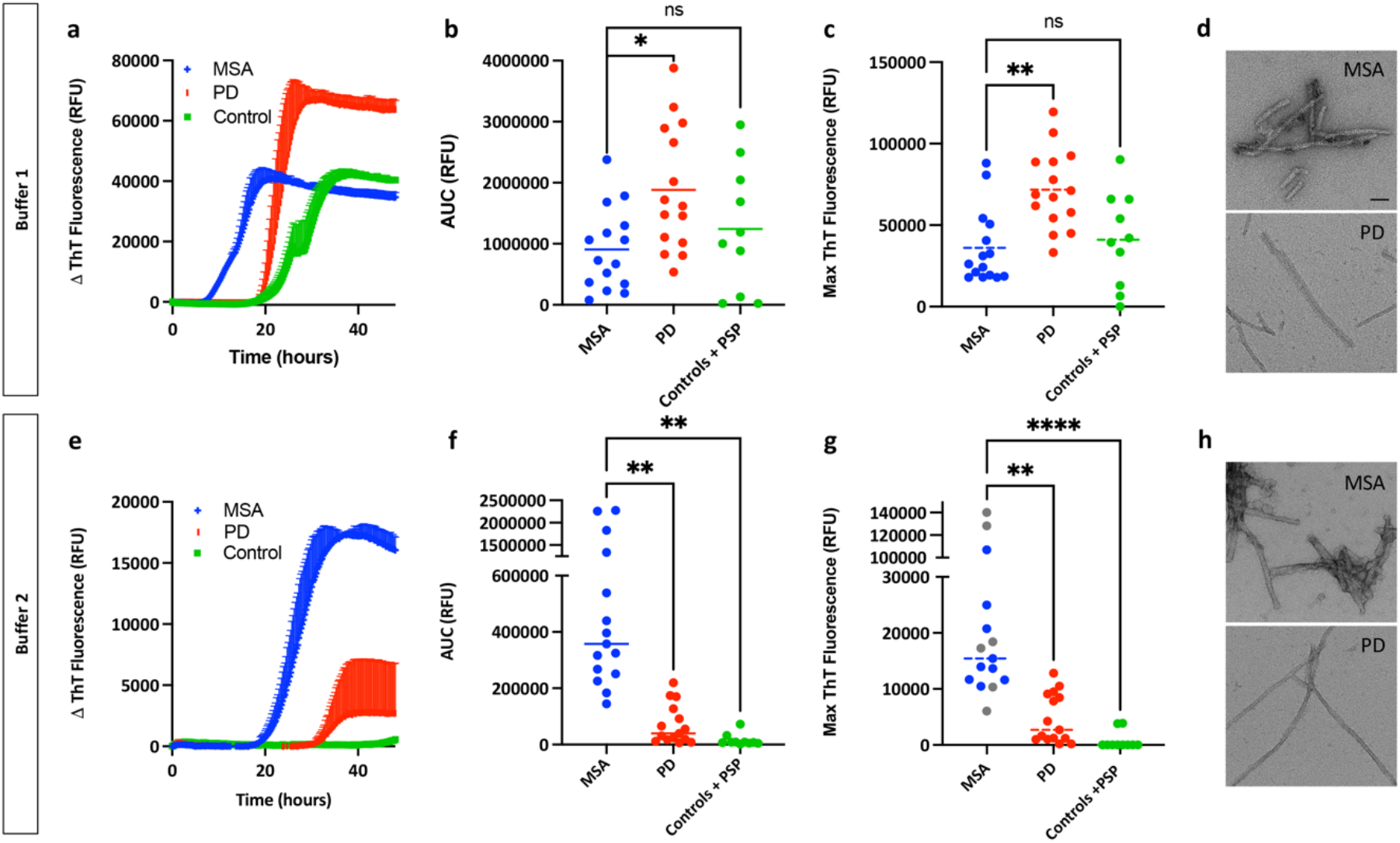
Interaction of ThT dye with α-synuclein aggregates derived from patients with PD or MSA is dependent on the RT-QuIC reaction buffer used. **a)** Representative aggregation curves (median case) of α-synuclein in the presence of brain homogenates from patients with PD (n = 15), patients with MSA (n = 15), patients with PSP (n=5) and controls (n = 5) amplified with the buffer 1. Data are mean ± s.e.m. of representative subjects measured in quadruplicate. Area under the curve values (**b**) and maximum ThT fluorescence values (**c**) for PD (n = 15), MSA (n = 15), PSP (n=5) and controls (n = 5). Each dot represents an individual biological sample measured in quadruplicate. **d)** At the ultrastructural level, RT-QuIC-derived MSA-derived fibrils are thicker and more twisted than the PD-derived fibrils, as determined by electron microscopy. **e)** Aggregation profiles of α-synuclein in the presence of brain homogenates from patients with PD (n = 15), patients with MSA (n = 15), patients with PSP (n=5) and controls (n = 5) amplified with the buffer 2. Data are mean ± s.e.m. of representative subjects measured in quadruplicate. Area under the curve values (**f**) and maximum ThT fluorescence values (**g**) for PD (n = 15), MSA (n = 15), PSP (n=5) and controls (n = 5). Each dot represents an individual biological sample measured in quadruplicate. The grey dots indicate the MSA patients that were selected for the brain regional analysis according to their divergent seeding capacity. **h)** At the ultrastructural level, RT-QuIC-derived MSA fibrils are thicker and twisted than the PD-derived fibrils, as determined by electron microscopy. \**P*<0.05, \*\**P*<0.01, \*\*\**P*<0.001, \*\*\*\**P* < 0.0001 by one-way analysis of variance (ANOVA) followed by Tukey’s multiple comparison test (b, c, f, g). Scale bar (d, h): 25 nm

### Characterization of the RT-QuIC end-products

To gain further insight regarding the structures of the aggregates that were amplified from patients with PD or with MSA, we analyzed the buffer 2 RT-QuIC end-products by CSA and thermolysin digestion. For these experiments, we examined the RT-QuIC end-products derived from the PD and MSA subjects used to seed the assays of the first phase of the study. As previously mentioned, under the CSA, the aggregates present in the MSA brain homogenate were significantly less stable following exposure to GdnCl than those found in the PD brain homogenate. This difference in conformational stability was preserved following the *in-vitro* amplification by RT-QuIC, and the end-products derived from the MSA brain were less stable than the PD ones (Fig. 4a-d). When the banding pattern of thermolysin-resistant α-synuclein species was examined, we found that the MSA homogenate and the MSA-derived RT-QuIC end-products possessed a band at ∼10 kDa and a prominent additional band at ∼15 kDa (Fig. 4e). However, when the PD samples were examined, the brain homogenate showed a band at ∼10 kDa, but the α-synuclein present in the PD RT-QuIC end-product was not thermolysin-resistant. These results, together with the ultrastructural data, show that, under our assay conditions, RT-QuIC reaction products seeded from MSA α-synuclein fibrils are conformationally distinguishable from RT-QuIC reaction products seeded from PD-derived fibrils.

**Fig. 4:**
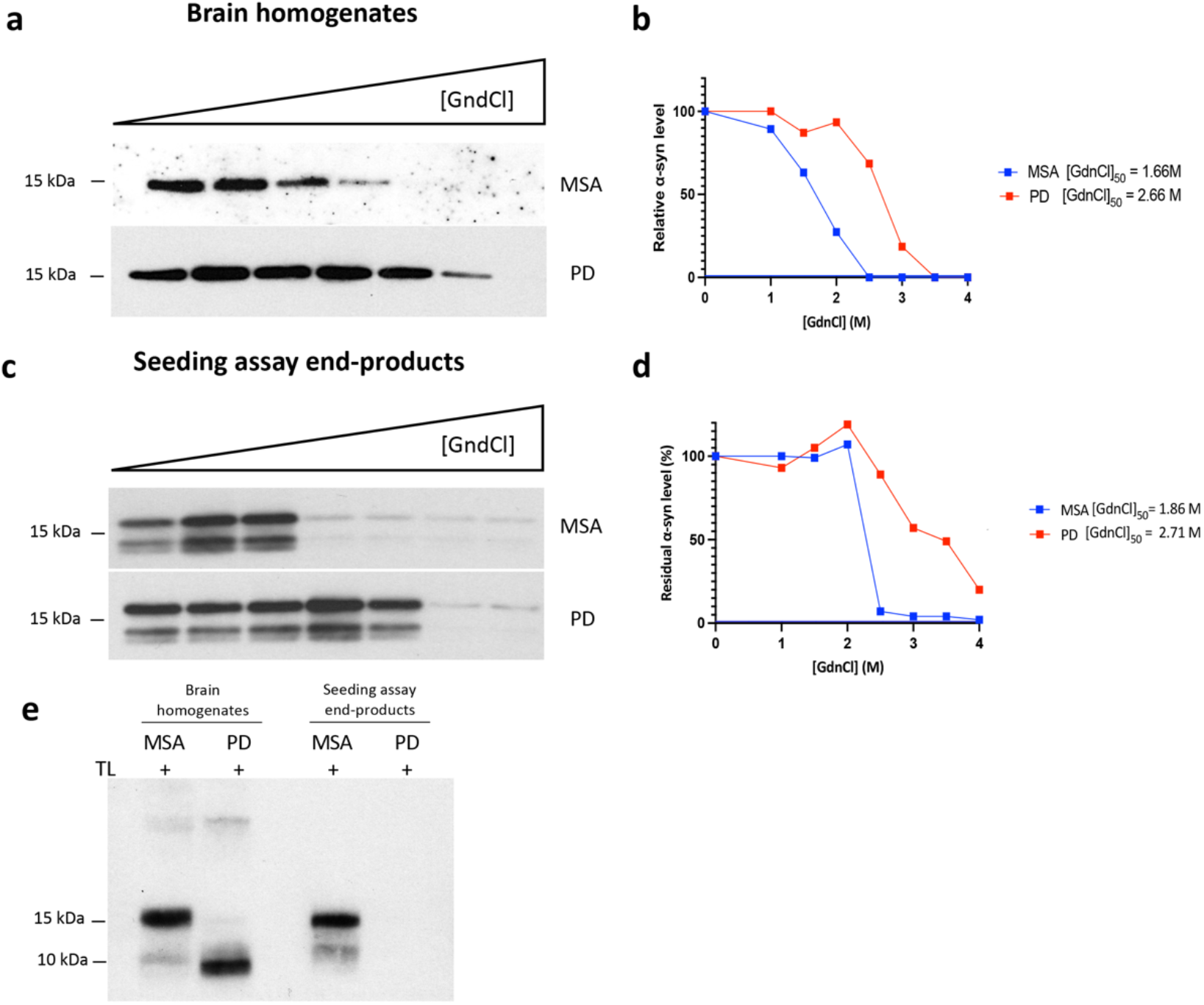
RT-QuIC reaction products seeded from MSA α-synuclein fibrils are conformationally distinguishable from RT-QuIC reaction products seeded from PD-derived fibrils. **a)** CSA for α-synuclein aggregates in brain extracts from patients with MSA and PD. Representative phosphorylated α-synuclein immunoblots and the resultant denaturation curves (**b**) are shown. The curves depict mean residual phosphorylated α-synuclein values following treatment with the indicated concentrations of GdnCl. Higher GdnCl_50_ values were obtained for α-synuclein aggregates in patients with PD, than for patients with MSA. **c**) CSA for the RT-QuIC-derived MSA and PD fibrils. Representative phosphorylated α-synuclein immunoblots and the resultant denaturation curves (**d**) are shown. MSA-derived fibrils are less stable than PD-derived fibrils. **e)** Immunoblots of detergent-insoluble α-synuclein species in brain homogenates from PD and MSA patients and their RT-QuIC-derived fibrils with digestion with thermolysin (TL).TL-resistant α-syn species were present in MSA and PD brain extracts, but TL-resistant α-syn species were only detectable in the MSA RT-QuIC-derived fibrils.

### MSA patients present distinct α-synuclein seeding

In addition to the observed differences in the AUC and ThT max values among the different groups of patients examined, when the individual values of the 15 MSA subjects were examined, we observed up to tenfold differences in AUC and ThT max values among MSA samples (Fig. 3f-g). Based on these data, the subjects with MSA included in our study were further subtyped into 3 groups: high, normal and low seeders, according to their ability to misfold monomers of recombinant α-synuclein *in vitro*. To test the hypothesis that heterogeneity in the α-synuclein seeding exists in MSA, from our initial cohort of 15 MSA patients, we selected 6 subjects according to their α-synuclein seeding (Fig. 3g) where 2 were classified as low seeders (subjects MSA 1 and MSA 6), 2 as normal seeders (subjects MSA 3 and MSA 4) and 2 as high seeders (subjects MSA 2 and MSA 5). The α-synuclein seeding behavior was then assessed in 13 different brain regions from these subjects. From each case, we microdissected and subsequently prepared a PBS-soluble extract from the following regions: anterior cingulate cortex, anterior cingulate white matter, frontal cortex, frontal white matter, putamen, globus pallidus, amygdala, hippocampus, temporal cortex, temporal white matter, substantia nigra, pons base and cerebellar white matter (Extended Data Fig. 2). All samples were run in quadruplicate, and the RT-QuIC curve obtained by each reaction was evaluated as a measure of the α-synuclein seeding behavior (Extended Data Fig. 3). The AUC values were converted into scores ranging from 0 to 3. With these values we generated a heatmap of α-synuclein seeding, to illustrate the differences found between brain regions and patients (Fig 5a). Interestingly, when the mean AUC values from all the regions included were analyzed, the two patients categorized as high seeders had significantly higher AUC values than the subjects categorized as normal (p<0.0001) or low (p<0.0001) seeders. When the different brain regions were examined individually, the pons base showed the highest α-synuclein seeding. Surprisingly, the cerebellar white matter and the putamen were the regions where the lowest α-synuclein seeding was observed. When the different cortices were examined, we found that the frontal and temporal cortex displayed a medium seeding, while the cingulate cortex had higher seeding. A higher seeding was consistently observed in the white matter of the three cortices when compared to the grey matter. Overall, the hippocampus displayed a medium seeding, while the α-synuclein present in globus pallidus, SN and the amygdala had a higher seeding. These data clearly demonstrate that 1) there is extensive heterogeneity in the α-synuclein seeding between different MSA patients, and 2) there is extensive heterogeneity in the seeding behavior of α-synuclein extracted from different brain regions within a given MSA patient.

**Fig. 5:**
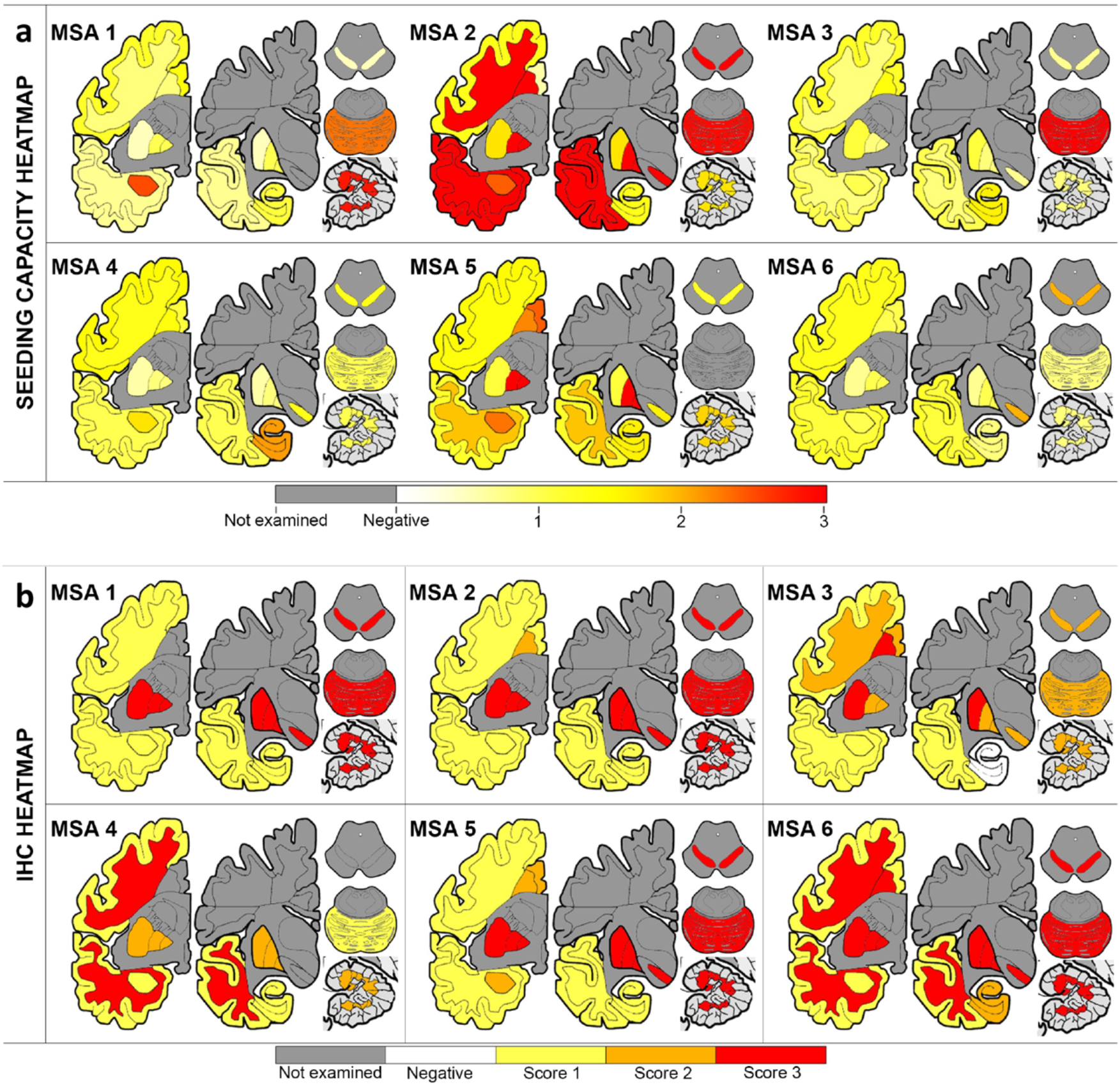
Extensive heterogeneity in α-synuclein seeding activity is observed in different MSA patients and between different brain regions. Heat mapping of α-synuclein seeding capacity (**a**) assessed by RT-QuIC and of aggregated α-synuclein (**b**) evaluated by immunohistochemistry using the conformational α-synuclein 5G4 antibody. The α-synuclein seeding capacity ranges from white (none) through yellow (low) and orange (medium) to red (high). The severity of α- synuclein pathology ranges from white (none) through yellow and orange to red (severe). Grey colored cortical regions indicate that the region was not evaluated.

### Biochemical and neuropathological characterization of α-synuclein

The factors that underlie the ability of α-synuclein from different patients with MSA to drive higher versus lower seeding are presently unknown. To investigate this further we conducted a biochemical examination of the α-synuclein derived from the brain homogenates coupled with a detailed immunohistochemical mapping of α-synuclein deposition in corresponding samples. First, we quantified the amount of total and aggregated α-synuclein using ELISAs and correlated these measures with their α-synuclein seeding behavior. MSA cases had 0.6–4 ng of total α-synuclein per mg of tissue in the PBS soluble fraction. No differences in the overall amount of α-synuclein were found between high, low and normal seeders. The regions where total α-synuclein was more abundant were the amygdala and the frontal cortex, whereas the cerebellar white matter and the pons were the regions where the α-synuclein was less expressed (Extended data Fig 4). The amount of aggregated α-synuclein was quantified using the α-synuclein Patho ELISA. MSA cases had 3–362 pg of aggregated α-synuclein per mg of tissue in the PBS soluble fraction. Interestingly, the amount of aggregated α-synuclein was significantly higher (p=0.0165) in the low seeders compared to the high seeders. The regions that had a higher burden of aggregated α-synuclein were the temporal and the anterior cingulate cortex, whereas, as occurred with the total α-synuclein levels, the cerebellar white matter and the pons were the regions where the least aggregated α-synuclein was found (Extended data Fig 5). The total amount of α-synuclein in the PBS-soluble fraction did not correlate with either the amount of aggregated α-synuclein (p=0.171) or with the α-synuclein seeding (p=0.648). However, the levels of aggregated α-synuclein were negatively correlated with the α-synuclein seeding (p=0.035, r= −0.241). These data, together with the striking seeding activity in soluble fractions from brain regions without noticeable accumulations of α-synuclein, lead us to test whether the heterogeneity between regions and patients was also maintained if larger, sarkosyl-insoluble (SI), α-synuclein aggregates were used to seed the RT-QuIC assay. To test this hypothesis, we extracted α-synuclein SI aggregates from the cerebellum, putamen and frontal cortex from one MSA case classified as a low seeder (MSA 1) and one MSA case categorized as a high seeder (MSA 2). Equal amounts (5μg) of total protein were used to seed all the reactions and quadruplicates from these samples together with their corresponding PBS-soluble fractions were evaluated. Furthermore, owing to the insoluble nature of the SI aggregates and possible inaccessibility of the fibril ends, we also included sonicated SI samples. When the RT-QuIC curves were evaluated, no significant differences between regions were observed in the SI aggregates, including those that were sonicated prior to the RT-QuIC. However, the SI aggregates from the three brain regions of the high seeder (MSA 2) aggregated faster and reached a higher fluorescence plateau than the SI aggregates from the low seeder (MSA 1). Regardless of the region or the patient examined, the SI aggregates promoted a faster aggregation and reached a higher fluorescence plateau than the α-synuclein aggregates present in the corresponding PBS-soluble fraction. The same pattern was observed when the differences in the α-synuclein aggregation kinetics were examined between SI aggregates with and without sonication, where the sonicated SI aggregates promoted a faster aggregation and reached a higher fluorescence plateau than the non-sonicated.

Finally, we aimed to evaluate whether the heterogeneity of α-synuclein seeding was reflected in immunohistological findings in the brains of the same subjects. First, we performed an epitope mapping to compare which α-synuclein antibody was able to detect the most MSA α-synuclein pathology in formalin-fixed paraffin-embedded tissue sections. Using consecutive sections from the putamen and the cerebellum from our cohort of 6 MSA patients, we performed morphometric analysis using four different α-synuclein antibodies: the 5G4 conformational antibody that recognizes aggregated forms of α-synuclein, the pSyn#64 that recognizes Ser129 phosphorylated α-synuclein, and two antibodies against the truncated (C-terminal cleaved) and nitrated forms of α-synuclein (clones A15127A and Syn514, respectively) (Extended data Fig 6). The conformational 5G4 antibody consistently exhibited more α-synuclein pathology in all the cases and regions examined followed by the two antibodies that recognized Ser 129 phosphorylated α-synuclein, and truncated α-synuclein (Extended data Fig 6m). Thus, using the 5G4 antibody, a semi-quantitative analysis of the α-synuclein aggregation burden was performed, the result of which are presented as a heatmap (Fig. 5b and Extended Data Fig. 7). The cerebellum and the putamen were the regions with the highest aggregated α-synuclein, with the SN, the pons base and the globus pallidus also exhibiting a high burden of aggregated α-synuclein. The grey matter from the temporal and frontal cortex were the regions where less amount of aggregated α-synuclein was found. In addition to the semi-quantitative analysis of the total burden of aggregated α-synuclein, the burden of GCIs and neuronal cytoplasmic inclusions (NCIs) was also examined (Extended Data Fig. 8). As expected, the presence of NCIs was less frequent than the presence of GCIs. The pons base, SN and putamen, followed by the hippocampus and the anterior cingulate cortex, were the regions where more NCIs were evident. When the GCIs burden was evaluated, the putamen and cerebellum, followed by the globus pallidus and the SN were the regions with more abundant GCIs, while the hippocampus and the frontal cortex has less abundant GCIs. Surprisingly, there was no significant difference in the burden of GCIs, NCIs or aggregated α-synuclein between high, low and normal seeders, and no correlation was found between the burden of GCIs (p=0.957) or NCIs (p=0.252) with the α-synuclein seeding.

## Discussion

Here we show that the physicochemical factors that govern the *in vitro* amplification of α-synuclein can be tailored to generate strain-specific reaction buffers for use in RT-QuIC. Using this novel approach, we have generated a streamlined RT-QuIC assay that 1) is able to measure the α-synuclein seeding of 96 MSA samples, run in quadruplicate, in less than 48 hours; 2) requires the use of a minimal amount of commercially available recombinant α-synuclein monomer (5μg); 3) requires the use of a minimal amount of brain material (5μg); 4) generates RT-QuIC-derived α-synuclein fibrils that are conformationally distinct between patients with different synucleinopathies; and 5) is capable of subtyping MSA brains according to their α-synuclein seeding behavior.

Using these assay conditions, we have conducted the first multi-regional evaluation of the α-synuclein seeding in MSA brains. MSA is clinically and pathologically heterogeneous, and although the distribution of GCIs and NCIs might correlate with the predominant clinical features^12,13^, the burden of α-synuclein inclusions does not fully explain the differences in clinical presentation and rate of disease progression exhibited in MSA. In recent years, mounting evidence highlights the important role of amyloidogenic proteins in soluble brain fractions to seed pathologic aggregation in a prion-like manner, that might contribute to clinical heterogeneity seen in patients with neurodegenerative diseases^23^. The capacity of misfolded α-synuclein to template monomeric α-synuclein has been exploited by several groups to generate an RT-QuIC assay able to measure seeding kinetics of α-synuclein in a range of samples in PD^17,20,24–26^. However, the reliable detection of α-synuclein seeding in MSA derived samples has been restricted to a single study^21^. To identify the optimal conditions to evaluate the α-synuclein seeding in MSA-derived samples, we performed a comprehensive analysis of 168 different reaction buffers, covering an array of pH and salts. We then validated the two conditions that conferred the optimal ability to discriminate between PD and MSA-derived samples in a larger cohort of neuropathologically confirmed cases. Although the same samples were used to seed the RT-QuIC reactions, the kinetics of the curves obtained using these two different buffers were significantly divergent, suggesting that the accessibility or the interaction of the ThT with α-synuclein from MSA and PD can be modulated by the type of salt, pH or ionic strength of the reaction buffer^21,22,27^. We selected buffer 2 as the optimal condition with which to evaluate α-synuclein seeding in MSA and further demonstrated that the RT-QuIC-derived MSA-fibrils maintained the biochemical properties of the MSA aggregates^28^ used to seed the reaction.

Our multi-region mapping of the α-synuclein seeding across 13 different brain regions in MSA revealed that the pons base had the highest seeding. Surprisingly, α-synuclein extracted from two of the most-affected regions in MSA, the cerebellum and the putamen, exhibited the lowest seeding observed. A possible explanation for these results is that the analysis of the α-synuclein seeding was performed using the PBS soluble fraction. It is plausible that the seeding activity of α-synuclein could diminish over time as the more soluble seeding competent α-synuclein species, present at earlier stages of disease, become sequestered in larger insoluble aggregates at later stages of the disease. This could then result in reduced seeding of α-synuclein extracted from regions affected earlier in the disease course^23,29^. This possibility led us to compare the seeding behavior of SI α-synuclein aggregates and the α-synuclein seeds present in the PBS soluble fraction in different brain regions. When the α-synuclein seeding was compared, regional differences were only observed using the PBS soluble fraction and not using the SI aggregates. However, regardless of the region or the patient examined, the SI aggregates consistently promoted a faster aggregation than the α-synuclein seeds present in the corresponding PBS-soluble fraction, suggesting that there are more competent α-synuclein species in the SI fraction. Our findings support earlier observations that modifications and solubility of α-synuclein in MSA may be more widespread than obvious histopathology^30,31^ and might be different between distinct synucleinopathies^31^. Future experiments will investigate the curious finding that PBS soluble α-synuclein is the major contributor to the observed regional differences in the seeding behavior in MSA. Interestingly, and irrespective of the differences found between the SI and the soluble fraction, striking differences in the seeding between patients were also observed.

Our study is subject to some limitations. Our data clearly demonstrate that the selection of the right conditions to conduct the amplification of the seeds is of major relevance. As a field, collectively our understanding of the optimal *in vitro* physicochemical factors with which specific seed conformers can propagate is still in its infancy. Using suboptimal conditions, an absence of seeding activity could reflect the absence of seeds, but also might reflect an inability of the specific RT-QuIC conditions to support the amplification of those particular seeds^32^.

Our findings pave the way for future studies to address the molecular mechanisms underlying the ability of α-synuclein from different patients with MSA to drive higher versus lower seeding. Indeed, our results support a model whereby the heterogeneity observed both between different brain regions and between different MSA patients might be due to differences in the cellular environment^6,8^ and to the presence of posttranslational modifications^9^ and other cofactors, such as p25α^33^, that may confer a selective pressure for one α-synuclein conformation over another^6^. Understanding the relationship between the seeding differences and biological activity of α-synuclein will be critical to developing personalized therapeutic strategies in the future. Furthermore, a deeper understanding of how physicochemical factors influence the aggregation of different α-synuclein strains will provide support for the development of vitally needed, rapid and sensitive *in vivo* assays for both the diagnosis and molecular subtyping of MSA and other synucleinopathies.

## Methods

### Human tissue samples

15 subjects with MSA, 15 subjects with PD, 5 controls and 5 subjects with PSP were selected from the from the University Health Network-Neurodegenerative Brain Collection (UHN-NBC, Toronto, Canada) and the Navarrabiomed Brain Bank (Pamplona, Spain) based on a definite neuropathological diagnostic. Age at death, sex and complete neuropathologically examination are listed in the Extended Data Table S1. Autopsy tissue from human brains were collected with informed consent of patients or their relatives and approval of local institutional review boards. Unfixed human brains were separated into 2 hemispheres, one of which was fixed in 10% formalin for 3 weeks, and regions of interest including 2.0 × 2.5 cm blocks of neocortical regions, hippocampus, amygdala, basal ganglia, thalamus, brainstem at different levels, and cerebellum were embedded in paraffin. 4-µm-thick paraffin-embedded tissue were cut and subsequently placed on slides (Superfrost, Thermo Scientific) for histological analysis. Prior to inclusion in the study, a systematic neuropathological examination was performed following diagnostic criteria of neurodegenerative conditions and co-pathologies^34^. The contralateral hemisphere was sliced coronally at the time of autopsy and immediately flash frozen and stored at −80 °C. Using a 3-mm biopsy punch, a microdissection of the following regions was performed: anterior cingulate cortex, anterior cingulate white matter, frontal cortex, frontal white matter, putamen, globus pallidus, amygdala, hippocampus, temporal cortex, temporal white matter, substantia nigra, pons base, and cerebellar white matter. All the punches were stored in low binding protein tubes (Eppendorf), immediately flash frozen and stored at −80 °C.

### Protein extraction

For the PBS-soluble fraction, 0.4-0.5 g of frozen microdissected tissue was thawed on wet ice and then immediately homogenized in 500 µl of PBS spiked with protease (Roche) and phosphatase inhibitors (Thermo Scientific) in a gentle-MACS Octo Dissociator (Miltenyi BioTec). The homogenate was transferred to a 1.5-ml low binding protein tube (Eppendorf) and centrifuged at 10,000g for 10 min at 4 °C. Then, the supernatant was collected and aliquoted in 0.5 mL low binding protein tubes (Eppendorf) to avoid excessive freeze–thaw cycles. Sarkosyl-insoluble material was extracted using 1-1.2g of frozen brain tissue from three brain regions (cerebellum, putamen and frontal cortex) of individuals with MSA, as previously described^9^. In brief, tissues were homogenized in 20 vol (v/w) extraction buffer consisting of 10 mM Tris-HCl, pH 7.5, 0.8 M NaCl, 10% sucrose and 1 mM EGTA. Homogenates were brought to 2% sarkosyl and incubated for 30 min, at 37 °C. Following a 10 min centrifugation at 10,000g, the supernatants were spun at 100,000g for 20 min. The pellets were resuspended in 500 µl/g extraction buffer and centrifuged at 3,000g for 5 min. The supernatants were diluted threefold in 50 mM Tris-HCl, pH 7.5, containing 0.15 M NaCl, 10% sucrose and 0.2% sarkosyl, and spun at 166,000g for 30 min. Sarkosyl-insoluble pellets were resuspended in 100 µl/g of 30 mM Tris-HCl, pH 7.4, and aliquoted in 0.5 mL low binding protein tubes (Eppendorf). For the detergent solubility assay, nine volumes of PBS brain homogenate were combined with a single volume of 10X detergent buffer (5% (v/v) Nonidet P-40, 5% (w/v) sodium deoxycholate, prepared in sterile PBS. The homogenate was centrifuged at 140,000g for 60 min at 4ºC, and the pellet was then resuspended in three volumes of ultra-pure water, and aliquoted in 0.5 mL low binding protein tubes (Eppendorf). A bicinchoninic acid protein (BCA) assay (Thermo Scientific) was performed to determine total protein concentration of all the aliquots.

### Histological analysis

Four µm-thick formalin-fixed paraffin-embedded tissue sections containing the thirteen anatomical regions selected for microdissection (see above) were examined. In addition to Hematoxylin and Eosin-Luxol Fast Blue, the following mouse monoclonal antibodies were used for immunohistochemistry: anti aggregated α-synuclein (5G4; 1:4000; Roboscreen), nitrated anti-α-synuclein (Syn514; 1:2000; Biolegend), C-terminal truncated x-122 anti-α-synuclein (A15127A; 1:2000; Biolegend) and anti-phospho-α-synuclein (pSyn#64; 1:10000; FUJIFILM Wako Pure Chemical Corporation). To map co-pathology, we used the following mouse monoclonal antibodies: anti-tau AT8 (pS202/pT205; 1:1000; Thermo Scientific), anti-phospho-TDP-43 (pS409/410; 1:2000; Cosmo Bio), and anti-Aβ (6F/3D; 1:50; Dako). Target retrieval for all antibodies, except for anti-Aβ and C-terminal truncated x-122 anti-α-synuclein, was performed using the DAKO EnVision FLEX Target Retrieval Solution. A 60 min incubation in 80% formic acid was used with the two antibodies where no EnVision FLEX antigen retrieval was performed. A second pretreatment of 1-and 5-min incubation in 80% formic acid was used for the phospho-TDP-43 and the 5G4 staining, respectively. The DAKO EnVision detection kit, peroxidase/DAB, rabbit/mouse (Dako) was used to visualize the antibody staining. For the comparison of different α-synuclein antibodies, immunostained sections against nitrated, truncated, phosphorylated, and aggregated (5G4) α-synuclein were scanned using Tissuescope™ and were cropped with the HuronViewer™ (Huron). Images were taken from the exact same location of the putamen and cerebellum across the different antibodies. Initially, 100 immunoreactive oligodendrocytes from each antibody, region and case with visible nucleus were optically dissected using Photoshop. Using Image J, the minimum and maximum areas (px^2^) of the 100 inclusions were recorded. The density of black dots per unit of inclusions were measured. Further, minimum and maximum values obtained from the previous step were used to measure the number of inclusions in each region and case for each antibody used. The amount of aggregated α-synuclein as well as the GCI and NCI burden were assessed using α-synuclein immunohistochemistry in the 13 brain regions above mentioned using the 5G4 staining. For semi-quantitative analyses, we used a 4-point scale:0, absent; 1, mild; 2, moderate; and 3, severe.

### ELISA

Human α-synuclein Patho and total ELISAs kits (Roboscreen GmbH) were used according to manufacturer’s protocol and as previously described^35^. Briefly, ELISA plates coated with their corresponding antibodies were used for incubation with diluted PBS-soluble brain homogenates. Controls and synthetic standards containing target antigens were incubated in parallel to brain samples for 24 hours at 2 – 10 °C, for the total α-synuclein kit and at room temperature with shaking (300 RPM) for the α-synuclein Patho kit. Plates were washed 5 times, and detection antibodies were added for 90 minutes at room temperature. After washing five times, the plates were stained using an ELISA staining kit (Roboscreen), and optical density was measured at 450/620 nm.

### SDS-PAGE and immunoblotting

Gel electrophoresis was performed using 4-12% or 12% Bolt Bis-Tris Plus gels (Thermo Scientific) for 35 min at 165 V. Proteins were transferred to 0.45 µm polyvinylidene fluoride membranes immersed in transfer buffer (25 mM Tris, pH 8.3, 0.192 M glycine, 20% (v/v) methanol) for 1 h at 35 V. Proteins were crosslinked to the membrane via 0.4% (v/v) paraformaldehyde incubation in PBS for 30 min at room temperature, with rocking. Membranes were blocked for 60 min at room temperature in blocking buffer (5% (w/v) skim milk in 1X TBST (TBS and 0.05% (v/v) Tween-20) and then incubated overnight at 4°C with primary antibodies diluted in blocking buffer. Primary antibodies: anti-Ser129-PαSyn EP1536Y (Abcam, ab51253; 1:4,000 dilution), anti-α-synuclein Syn-1 (BD Biosciences, 610786; 1:10,000 dilution). Membranes were then washed 3x with TBST and then incubated, for 60 min at room temperature, with horseradish peroxidase-conjugated secondary antibodies (Bio-Rad, 172-1019 or 172-1011) diluted 1:10,000 in blocking buffer. Following another 3x washes with TBST, immunoblots were developed using Western Lightning enhanced chemiluminescence (ECL) Pro (PerkinElmer) and imaged using X-ray film or the LiCor Odyssey Fc system.

### Conformational stability assays of brain-derived α-synuclein aggregates

20µL of 2X guanidine hydrochloride (GdnCl) stocks were added to an equal volume of detergent-extracted brain homogenates to yield final GdnCl concentrations of 0, 1, 1.5, 2, 2.5, 3, 3.5 and 4 M. PBS-soluble brain samples were incubated at room temperature with shaking for 120 min (800 RPM) before being diluted to 0.4 M GdnCl in PBS containing 0.5% (w/v) sodium deoxycholate and 0.5% (v/v) NP-40. Following high speed ultracentrifugation at 100,000 x g for 60 min at 4ºC pellets were resuspended in 1x LDS loading buffer and boiled for 10 min. Levels of residual α-synuclein were determined by SDS–PAGE followed by immunoblotting, as above mentioned. Densitometry was performed using ImageJ, and values were normalized to the sample with the highest intensity, which was set at 100%. GdnCl_50_ values, the concentration of GdnCl at which 50% of the aggregates are solubilized, were determined by nonlinear regression using the sigmoidal dose–response (variable slope) equation in GraphPad Prism, with the top and bottom values fixed at 100 and 0, respectively.

### Thermolysin (TL) digestion of brain-derived α-synuclein aggregates

Detergent-extracted brain homogenate was diluted into 1X detergent buffer [0.5% (v/v) Nonidet P-40, 0.5% (w/v) sodium deoxycholate in PBS] containing TL at a concentration of 50 µg/mL. Samples were incubated at 37°C with continuous shaking (600 RPM) for 1 hr. Digestions were halted with the addition of EDTA to final concentration of 2.5 mM, and samples were ultracentrifuged at 100,00 x g for 60 min at 4°C. Supernatant was discarded and pellets resuspended via boiling in 1X LDS buffer containing 2.5% β-mercaptoethanol. Samples were then analyzed via SDS-PAGE followed by immunoblotting, as described above.

### RT-QuIC assay and buffer preparation

RT-QuIC reactions were performed in white 384-well plates with a clear bottom (Nunc). Recombinant α-synuclein (rPeptide) was thawed from −80 ◦C storage, reconstituted in HPLC-grade water (Sigma) and filtered through a 100-kDa spin filter (Thermo Scientific) in 500-µL increments. All the reagents used for the reaction buffers were purchased from Sigma. 10 µL of the biological sample (5 µg of total protein from freshly thaw brain homogenates diluted in PBS) was added to wells containing 20µL of the reaction buffer, 10µL of 50µM Thioflavin T and 10µL of 0.5mg/ml of monomeric recombinant α-synuclein. 1 µg of recombinant α-synuclein fibrils was used as positive control, while 5 µg of total protein from the frontal cortex of control subjects was used as a negative control for the RT-QuIC reactions. The plate was sealed and incubated at 37°C in a BMG FLUOstar Omega plate reader with cycles of 1 min shaking (400 rpm double orbital) and 14 min rest for a period of 72 hours. ThT fluorescence measurements (450+/-10 nm excitation and 480+/-10nm emission, bottom read) was taken every 15 min for 72 hours.

### Conformational stability assays of RT-QuIC-derived α-synuclein fibrils

20µL of 2X GdnCl stocks were added to an equal volume of RT-QuIC-derived α-synuclein fibrils to yield final GdnCl concentrations of 0, 1, 1.5, 2, 2.5, 3, 3.5 and 4 M. RT-QuIC-derived α-synuclein fibrils were incubated at room temperature with shaking for 2 hours (800 RPM) before being diluted to 0.4 M GdnCl in PBS containing 0.5% (w/v) sodium deoxycholate and 0.5% (v/v) NP-40. Following high speed ultracentrifugation at 100,000 x g for 60 min at 4 ºC pellets were resuspended in 1x LDS loading buffer and boiled for 10 minutes. Levels of residual α-synuclein were determined by SDS–PAGE, as previously described, followed by immunoblotting. Densitometry was performed using ImageJ, and values were normalized to the sample with the highest intensity, which was set at 100%. GdnCl_50_ values, the concentration of GdnCl at which 50% of the aggregates are solubilized, were determined by nonlinear regression using the sigmoidal dose–response (variable slope) equation in GraphPad Prism, with the top and bottom values fixed at 100 and 0, respectively.

### Thermolysin digestion of RT-QuIC-derived α-synuclein fibrils

RT-QuIC-derived α-synuclein fibrils were diluted into 1X detergent buffer (0.5% (v/v) Nonidet P-40, 0.5% (w/v) sodium deoxycholate in PBS) containing TL at a concentration of 5 µg/mL. Samples were incubated at 37°C with continuous shaking (600 RPM) for 60 min. Digestions were halted with the addition of ethylenediamine tetraacetic acid (EDTA) to final concentration of 2.5 mM, and samples were ultracentrifuged at 100,00 x g for 60 min at 4°C. Supernatant was discarded and pellets resuspended via boiling in 1X LDS buffer containing 2.5% β-mercaptoethanol. Samples were then analyzed via SDS-PAGE followed by immunoblotting, as described above.

### Electron microscopy

Negative-stain electron microscopy was used to examine the ultrastructural characteristics of the MSA and PD RT-QuIC-derived fibrils. Aliquots (5 µl of α-synuclein fibril preparations) containing the RT-QuIC-derived α-synuclein fibrils were loaded onto freshly glow-discharged 400 mesh carbon-coated copper grids (Electron Microscopy Sciences) and adsorbed for 1 min. Grids were then washed with 50 µl each of 0.1 M and 0.01 M ammonium acetate and stained with 2 × 50 µl of freshly filtered 2% uranyl acetate. Once dry, grids were visualized using a Talos L120C transmission electron microscope (Thermo Fisher) using an acceleration voltage of 200 kV. Electron micrographs were recorded using an Eagle 4kx4k CETA CMOS camera (Thermo Fisher).

### Statistics and reproducibility

Statistical analyses were performed using GraphPad Prism (v.9) with a significance threshold of *P* = 0.05. RT-QuIC relative fluorescence responses were also analyzed and plotted using the software GraphPad Prism (v.9). The area under the curve (AUC), the maximum intensity of fluorescence (ThT max), and the lag phase were extracted, and the normality of the distribution of these variables from each group was assessed using the Kolmogorov–Smirnov test. The Data were compared using unpaired two-tailed t-tests and one-way ANOVA with Tukey’s multiple comparisons test. Two-tailed Spearman *r* non-parametric correlations were used to correlate different variables obtained from single individuals using SPSS. The Spearman correlation coefficient *r* and the *P* value are indicated. Data collection and analyses were not performed blinded to the conditions of the experiments. All experiments on RT-QuIC-derived α-synuclein fibrils were performed using a minimum of three independent fibril preparations per condition. For RT-QuIC analysis, all the brain regions were analyzed in quadruplicate. For the CSA and the thermolysin digestion, the same MSA and PD brains were analyzed. For ELISAs, all the samples were measured in triplicates. For the electron microscopy analysis, 20 electron micrographs from 3 MSA and 3 PD subjects were analyzed, and a similar ultrastructure was obtained for every sample within a group.

### Reporting Summary

Further information on research design will be available in the Nature Research Reporting Summary linked to this article.

## Data availability

The data that support the findings of this study are available from the corresponding author upon reasonable request.

## Acknowledgements

This study was supported by the Edmond J Safra Philanthropic Foundation and Rossy Foundation.

## Author contributions

Designed the experiments: IMV, GGK

Conducted the experiments and provided the samples: IMV, AK, HHCL, RWLS, SA, AG, MRL, MS, GGK

Analyzed and interpreted the data: IMV, NPV, HHCL, RWLS, JCW, AEL, GGK Wrote the manuscript: IMV, NPV, GGK

AEL and JCW provided critical input on the manuscript

All authors edited and approved the manuscript

