## Extended files for "A novel approach to evaluate alpha-synuclein seeding shows a wide heterogeneity in multiple system atrophy"

### SUPPLEMENTAL FILES

Ivan Martinez-Valbuena[1], Naomi P. Visanji[2,3,4], Ain Kim[1], Heather H. C. Lau[1,5], Raphaella W. L. So[1,5], Sohaila Alshimemeri[2], Andrew Gao[3,6], Michael Seidman[3,6], Maria R. Luquin[7], Joel C. Watts[1,5], Anthony E. Lang[1,2,4] and Gabor G. Kovacs[1,2,3,4,6*]

1. Tanz Centre for Research in Neurodegenerative Disease, University of Toronto, Toronto, Ontario, Canada
2. Edmond J. Safra Program in Parkinson's Disease and the Morton and Gloria Shulman Movement Disorders Clinic, Toronto Western Hospital, Toronto, Ontario, Canada.
3. Department of Laboratory Medicine and Pathobiology, University of Toronto, Toronto, Ontario, Canada
4. Krembil Brain Institute, University Health Network, Toronto, Ontario, Canada
5. Department of Biochemistry, University of Toronto, Toronto, Ontario, Canada
6. Laboratory Medicine Program, University Health Network, Toronto, Ontario, Canada
7. Department of Neurology, Clinica Universidad de Navarra, Pamplona, Navarra, Spain

*

**Abstract: 150 words**

| | |
|---|---|
| Word count title: | 16 |
| Word count abstract: | 142 |
| Word count text: | 4526 |
| Number of figures: | 5 |
| Supplemental files: | 9 supplementary figures + 1 table in the extended data |

**Corresponding author:** Gabor G. Kovacs MD PhD FRCPC

University of Toronto, Tanz Centre for Research in Neurodegenerative Disease (CRND), Krembil Discovery Tower, 60 Leonard Ave Toronto On, M5T 0S8, Canada; Tel: +1 (416) 507-6858;.

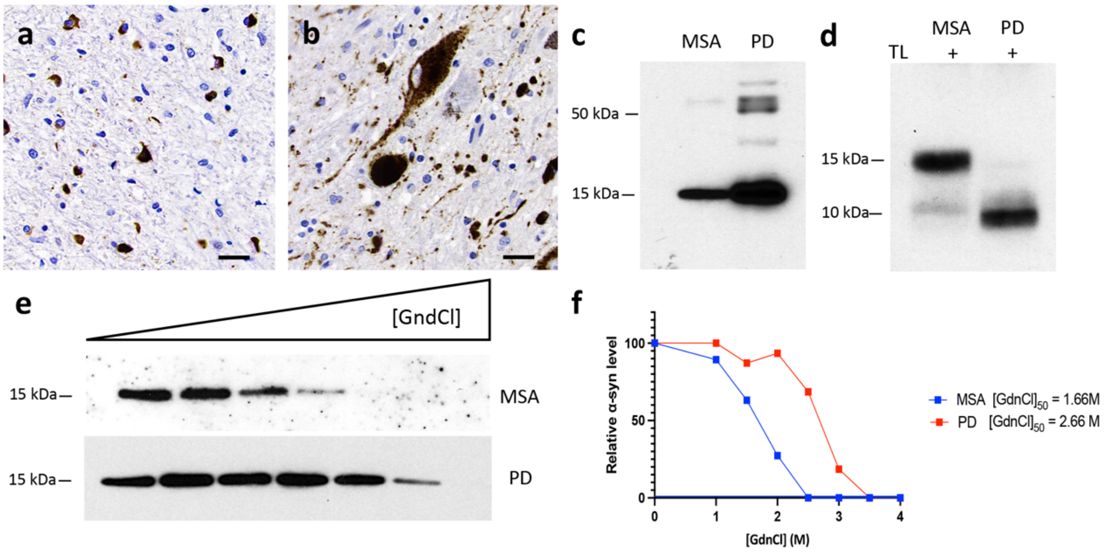


**Extended data Fig.1: Conformational discrimination of α-synuclein strains in PD and MSA brain. a)** Representative immunohistochemistry image for aggregated α-synuclein in the cerebellum white matter from a MSA patient showing GCIs. **b)** Representative immunohistochemistry image for aggregated α-synuclein in the SN from a PD patient showing Lewy bodies and Lewy neurites. **c)** Representative α-synuclein immunoblots showing the different molecular weights of the α-synuclein species present in the brain homogenates of the PD and the MSA patients after performing a detergent insolubility assay. **d)** Thermolysin digestion of brain extracts from the cerebellum and the SN of a MSA and PD patient, respectively. **e)** CSA for α-synuclein aggregates in brain extracts from patients with MSA or PD. Representative α-synuclein immunoblots and the resultant denaturation curves **(f)** are shown. The curves depict mean residual α-synuclein values following treatment with the indicated concentrations of GdnCl. Higher $GdnCl_{50}$ values were obtained for α-synuclein aggregates in the patient with PD than for the patient with MSA**.**

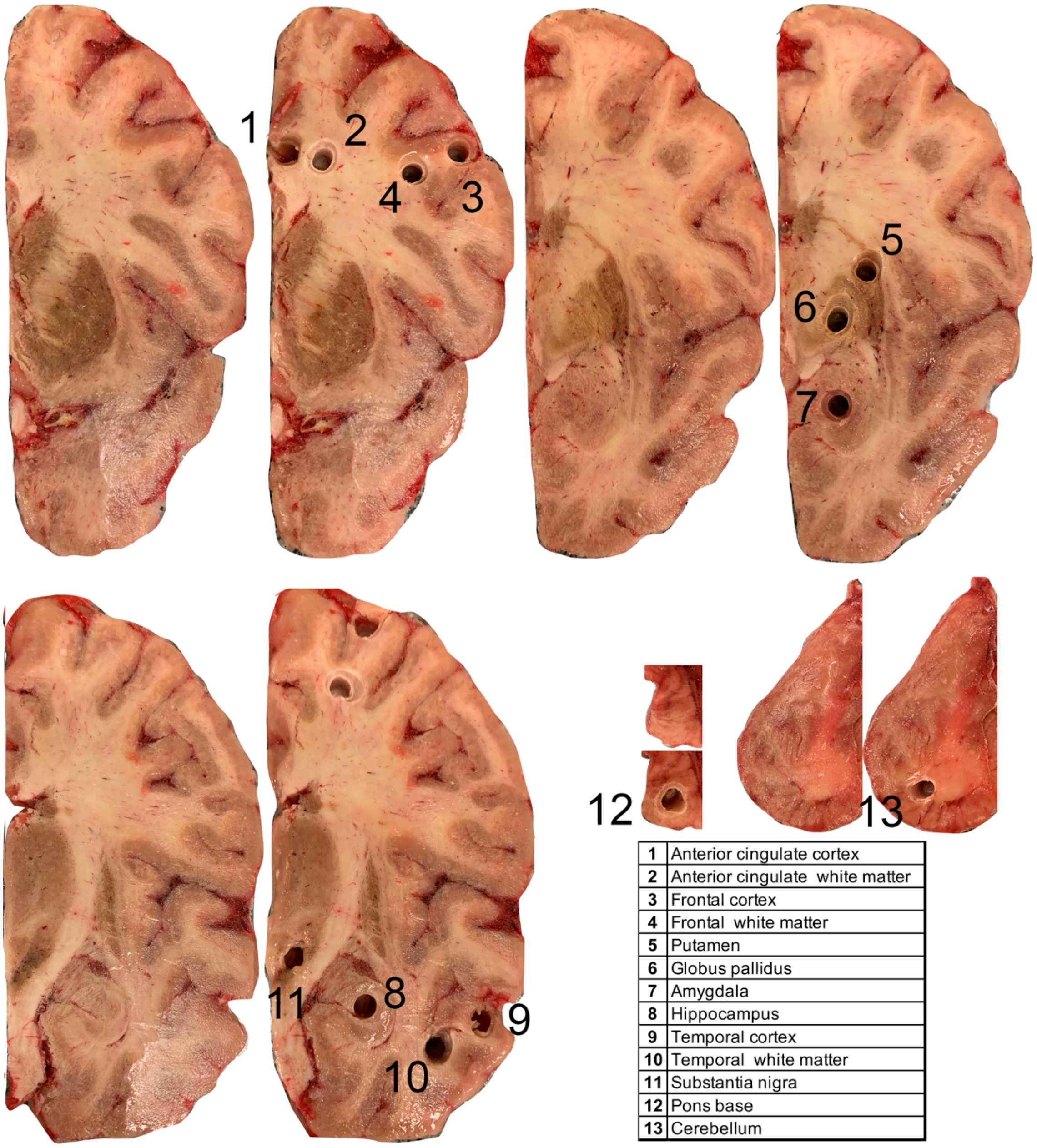


| | |
|---|---|
| 1 | Anterior cingulate cortex |
| 2 | Anterior cingulate white matter |
| 3 | Frontal cortex |
| 4 | Frontal white matter |
| 5 | Putamen |
| 6 | Globus pallidus |
| 7 | Amygdala |
| 8 | Hippocampus |
| 9 | Temporal cortex |
| 10 | Temporal white matter |
| 11 | Substantia nigra |
| 12 | Pons base |
| 13 | Cerebellum |

**Extended data Fig.2: Dissection of brain regions for protein extraction and α-synuclein seeding capacity evaluation.** Coronal sections from frozen brains were used to dissect, up to 13 different brain regions in all the subjects included in the study, using a 3-mm punch biopsy tool. Different biopsy punch tools were used between subjects and regions to avoid cross-contamination. Once the brain regions were dissected, they were stored at -80ºC until the protein was extracted.

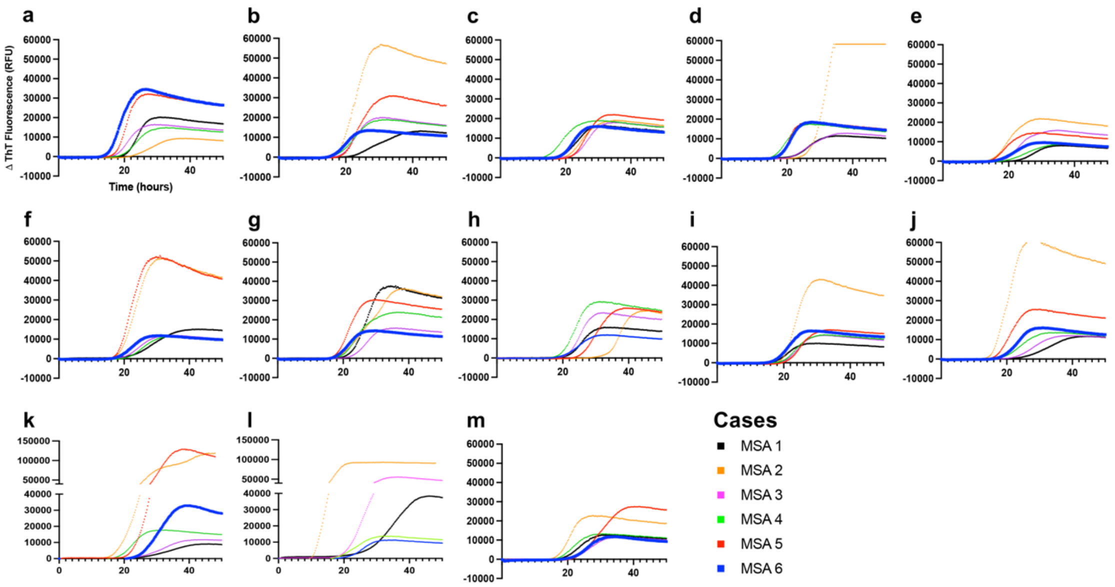


**Extended data Fig.3: The inter-individual α-synuclein seeding capacity and intra-individual α-synuclein seeding capacity is distinct between MSA patients and brain regions.** Kinetic curves of α-synuclein seeding activity measured by RT- QuIC in **a)** anterior cingulate cortex **b)** anterior cingulate white matter **c)** frontal cortex **d)** frontal white matter **e)** putamen **f)** globus pallidus **g)** amygdala **h)** hippocampus **i)** temporal cortex **j)** temporal white matter **k)** substantia nigra **l)** pons base and **m)** cerebellum white matter from 6 different MSA patients. Each curve depicts the average of quadruplicates. Standard deviation (SD) was hidden to make the image more readable.

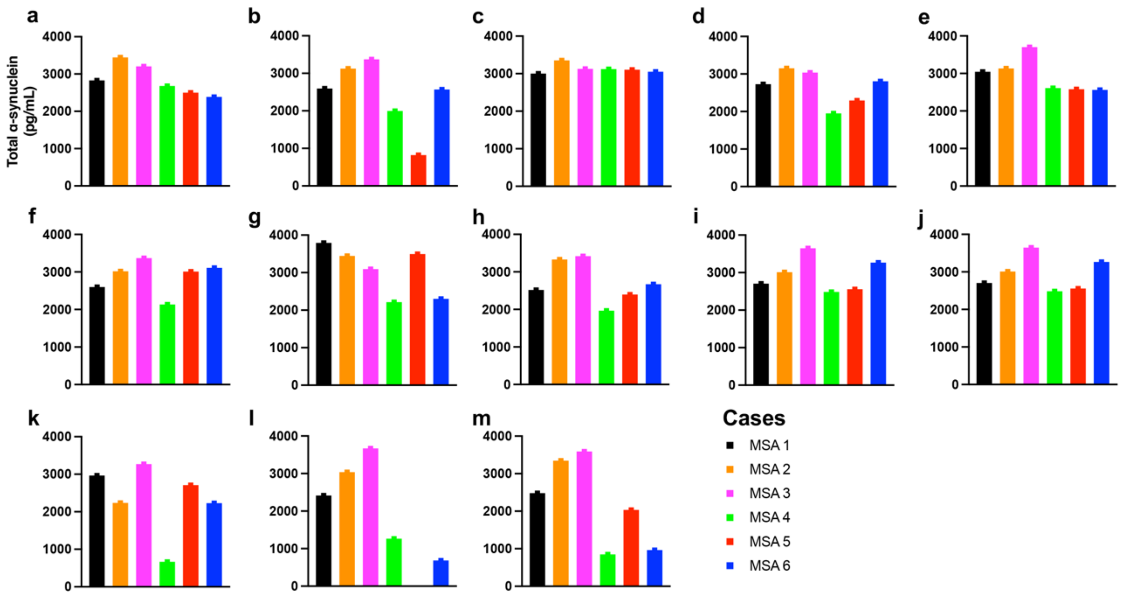


**Extended data Fig.4: The levels of total α-synuclein in each brain region are relatively uniform between MSA patients.** The amount of total α-synuclein was quantified, using an ELISA assay, in **a)** anterior cingulate cortex **b)** anterior cingulate white matter **c)** frontal cortex **d)** frontal white matter **e)** putamen **f)** globus pallidus **g)** amygdala **h)** hippocampus **i)** temporal cortex **j)** temporal white matter **k)** substantia nigra **l)** pons base and **m)** cerebellum white matter from 6 different MSA patients.

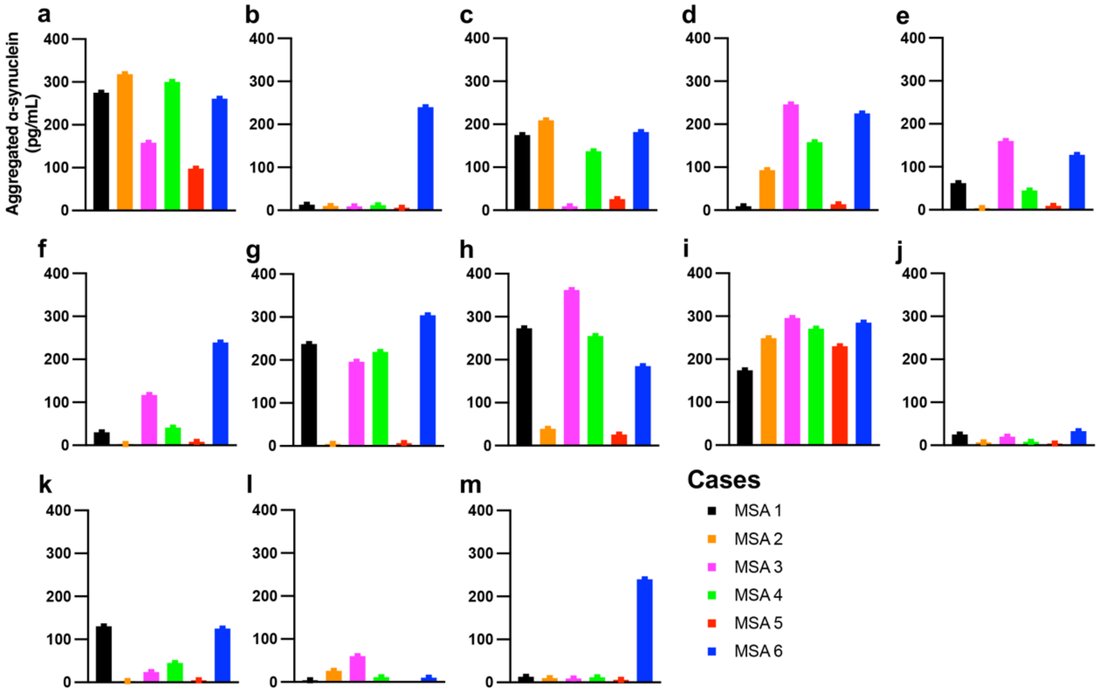


**Extended data Fig.5: The burden of aggregated α-synuclein varies across patients and brain regions in MSA.** The amount of aggregated α-synuclein was quantified using the α-synuclein Patho ELISA assay, that uses the 5G4 antibody as capture antibody, in **a)** anterior cingulate cortex **b)** anterior cingulate white matter **c)** frontal cortex **d)** frontal white matter **e)** putamen **f)** globus pallidus **g)** amygdala **h)** hippocampus **i)** temporal cortex **j)** temporal white matter **k)** substantia nigra **l)** pons base and **m)** cerebellum white matter from 6 different MSA patients

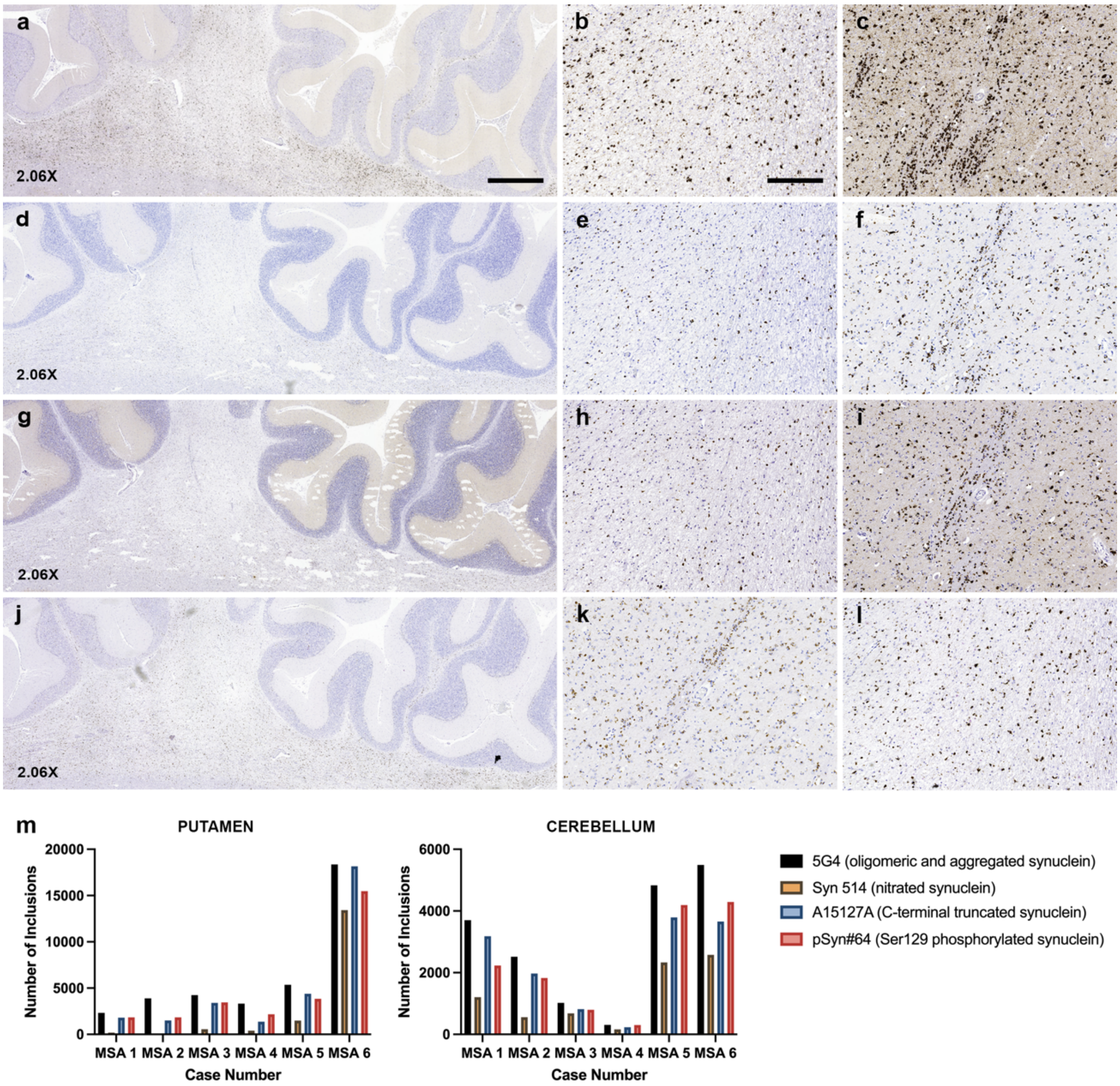


**Extended Data Fig. 6: The extent of pathology detected by different** a-synuclein **antibodies is not uniform.** Representative images showing **a)** 5G4 immunoreactivity in the cerebellum of MSA 6 at low magnification (2.06X); **b-c)** 5G4 immunoreactivity in the cerebellum and putamen of MSA 6 at higher magnification (6.25X), respectively; **d)** Ser 129 phosphorylated alpha-synuclein immunoreactivity in the cerebellum of MSA 6 at low magnification (2.06X); **e-f)** phosphorylated alpha-synuclein inclusions in the cerebellum and putamen of MSA 6 at higher magnification (6.25X), respectively; **g)** immunoreactivity showing nitrated alpha-synuclein inclusions in the cerebellum of MSA 6 at low magnification (2.06X); **h-i)** nitrated alpha-synuclein inclusions in the cerebellum and putamen of MSA 6 at higher magnification (6.25X), respectively; **j)** immunoreactivity showing truncated alpha-synuclein inclusions in the cerebellum of MSA 6 captured at low magnification (2.06X); **k-l)** truncated alpha-synuclein inclusions in the cerebellum and putamen of MSA 6 at higher magnification (6.25X), respectively. **m)** number of inclusions observed using the 4 different antibodies against α-synuclein in the putamen and cerebellum of 6 MSA patients. Scanned images were cropped at the same region to match the location in the images taken from different alpha-synuclein antibodies at the same magnification and size. Scale bar in a represents 4000 μm in a, d, g, j and scale bar represents 125 μm in b, c, e, f, h, i, k, l.

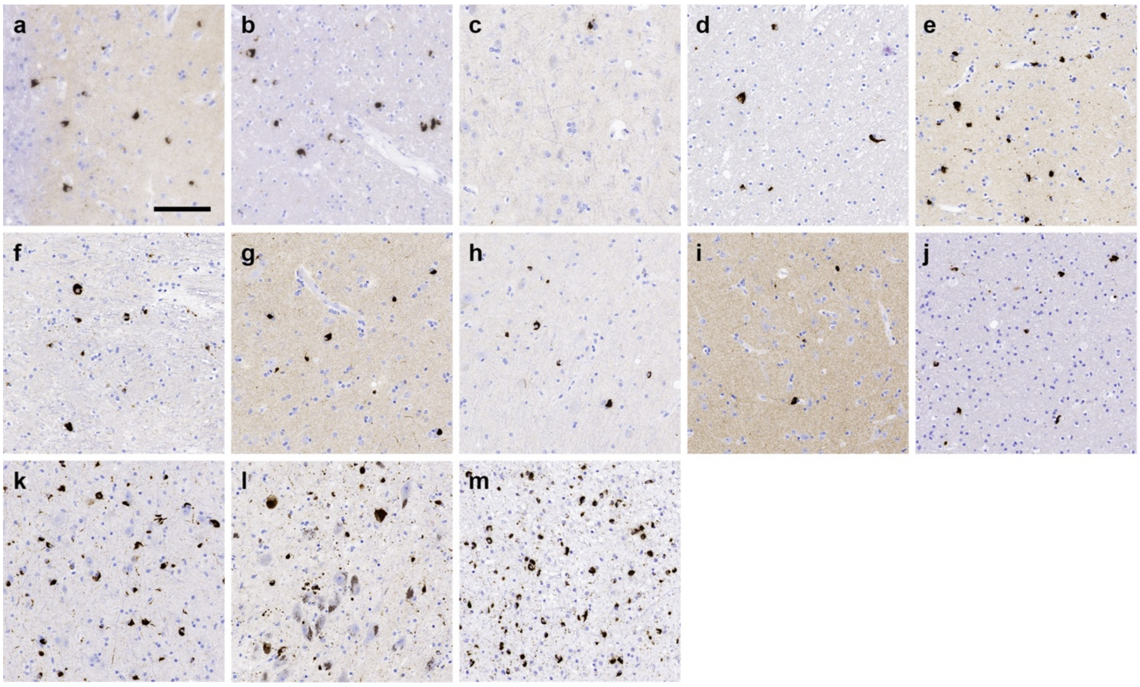


**Extended data Fig.7: GCIs and NCIs deposition across different brain regions in MSA patients.** Representative immunohistochemistry images for aggregated α-synuclein in **a)** anterior cingulate cortex **b)** anterior cingulate white matter **c)** frontal cortex **d)** frontal white matter **e)** putamen **f)** globus pallidus **g)** amygdala **h)** hippocampus **i)** temporal cortex **j)** temporal white matter **k)** substantia nigra **l)** pons base and **m)** cerebellum white matter from the MSA5 patient. Scale bar in '**a**'= 50 µm (applies to all images).

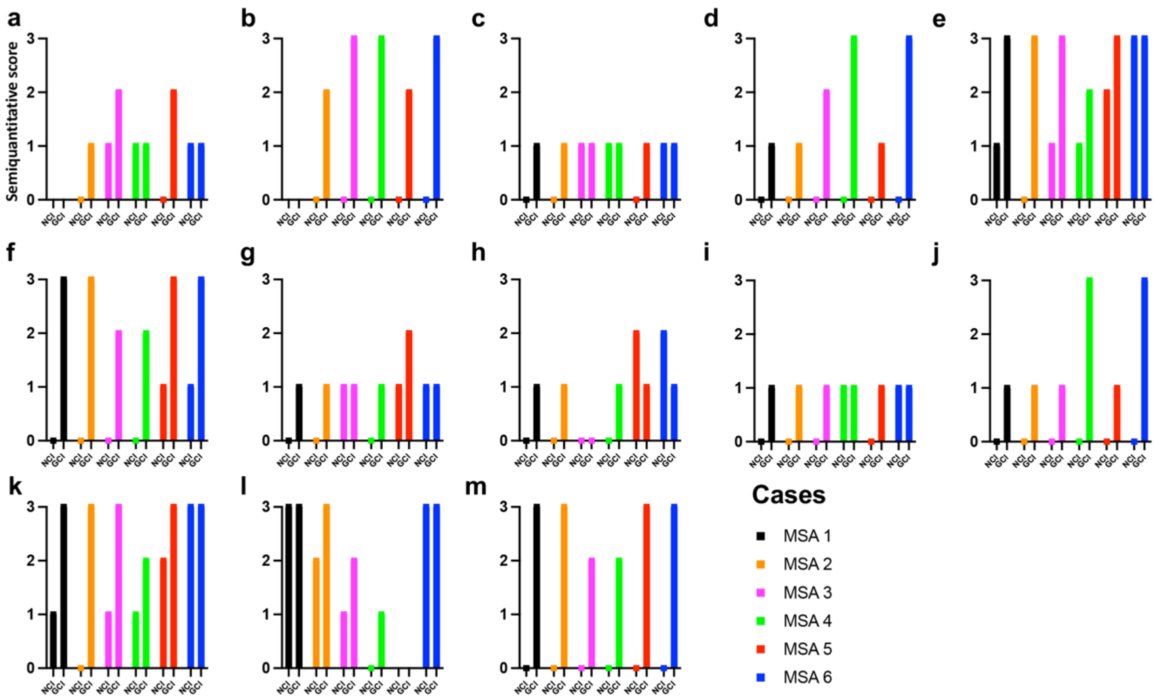


**Extended data Fig. 8: The burden of GCIs and NCIs varies across different brain regions in MSA.** The burden of GCIs (second bar) and NCIs (first bar) was evaluated using a semiquantitative 4-point scale where 0 was the absence of α-synuclein pathology; 1 was mild; 2 was moderate and 3 was severe. The evaluation was performed in the **a)** anterior cingulate cortex **b)** anterior cingulate white matter **c)** frontal cortex **d)** frontal white matter **e)** putamen **f)** globus pallidus **g)** amygdala **h)** hippocampus **i)** temporal cortex **j)** temporal white matter **k)** substantia nigra **l)** pons base and **m)** cerebellum white matter from 6 different MSA patients.

**Extended data Table 1. Co-pathology for Case Selection**

| Case | Code | Source | Sex | Age | NIA-AA | Cerebral Amyloid Angiopathy | Lewy Body Pathology | Lewy Body Braak Stage | LATE-NC Stage | Other Pathologic Diagnosis |
|---|---|---|---|---|---|---|---|---|---|---|
| **LBD 1** | 561 | Canada | F | 59 | A3B3C3 | - | Neocortical (diffuse) | Stage 5 | - | - |
| **LBD 2** | 973 | Canada | F | 82 | A2B3C3 | Aβ-positive Type 2 | Neocortical (diffuse) | Stage 5 | Stage 2 | - |
| **LBD 3** | 1074 | Canada | M | 73 | A2B3C2 | Aβ-positive Type 2 | Neocortical (diffuse) | Stage 5 | Stage 2 | - |
| **LBD 4** | 1075 | Canada | M | 71 | A2B2C3 | - | Neocortical (diffuse) | Stage 5 | - | - |
| **LBD 5** | 1153 | Canada | M | 79 | A3B3C3 | Aβ-positive Type 1 and 2 | Neocortical (diffuse) | Stage 5 | - | - |
| **LBD 6** | 1160 | Canada | M | 84 | A2B2C2 | Aβ-positive Type 1 | Limbic (transitional) | Stage 4 | Not Assessed | - |
| **LBD 7** | 1181 | Canada | F | 78 | A3B3C3 | - | Neocortical (diffuse) | Stage 5 | Stage 2 | - |
| **LBD 8** | 1195 | Canada | M | 80 | A2B3C2 | Aβ-positive Type 1 and 2 | Limbic (transitional) | Stage 4 | Stage 1 | - |
| **LBD 9** | 1245 | Canada | F | 82 | A3B3C3 | Aβ-positive Type 2 | Limbic (transitional) | Stage 4 | Stage 2 | - |
| **LBD 10** | 1239 | Canada | M | 73 | A2B2C2 | - | Neocortical (diffuse) | Stage 5 | - | - |
| **LBD 11** | 1265 | Canada | F | 88 | A2B2C2 | Aβ-positive Type 1 and 2 | Limbic (transitional) | Stage 4 | Stage 2 | - |
| **LBD 12** | 1305 | Canada | F | 94 | A2B2C2 | Aβ-positive Type 2 | Neocortical (diffuse) | Stage 5 | - | - |
| **LBD 13** | 1404 | Canada | F | 73 | A2B3C2 | - | Limbic (transitional) | Stage 4 | Stage 2 | - |
| **LBD 14** | 1409 | Canada | F | 76 | A3B3C3 | Aβ-positive Type 2 | Neocortical (diffuse) | Stage 5 | Stage 2 | - |
| **LBD 15** | 1424 | Canada | M | 69 | A1B1C0 | - | Limbic (transitional) | Stage 4 | - | |

| Case | Code | Source | Sex | Age | NIA-AA | Cerebral Amyloid Angiopathy | Lewy Body Pathology | Lewy Body Braak Stage | LATE-NC Stage | Other Pathologic Diagnosis |
|---|---|---|---|---|---|---|---|---|---|---|
| **MSA 1** | 585 | Canada | F | 68 | A1B1C0 | Aβ-positive Type 2 | - | - | - | ARTAG Medial Temporal Lobe WM |
| **MSA 2** | 1381 | Canada | M | 72 | A1B0C0 | - | - | - | - | Hemorrhage: Pons Base & Thalamus |
| **MSA 3** | 1461 | Canada | F | 76 | A0B1C0 | - | - | - | - | PART (Braak stage II) |
| **MSA 4** | R-NBC-20-31 | Canada | M | 61 | A0B1C0 | - | - | - | - | PART (Braak stage II) |
| **MSA 5** | R-NBC-20-9 | Canada | M | 64 | A0B1C0 | - | - | - | - | PART (Braak stage II), AGD (Stage II) |
| **MSA 6** | R-NBC-20-19 | Canada | M | 62 | A0B1C1 | - | - | - | - | ARTAG Medial Temporal Lobe GM |
| **MSA 7** | 1613 | Canada | M | 62 | A0B1C0 | - | - | - | - | - |
| **MSA 8** | 1704 | Canada | M | 73 | A0B1C0 | - | - | - | - | - |
| **MSA 9** | A-14-02 | Canada | M | 71 | A0B1C0 | - | - | - | - | - |
| **MSA 10** | BCN-306 | Spain | F | 65 | A0B1C0 | - | - | - | - | - |
| **MSA 11** | BCN-321 | Spain | M | 77 | A0B1C0 | - | - | - | - | - |
| **MSA 12** | BCN-371 | Spain | F | 80 | A0B1C0 | - | - | - | - | - |
| **MSA 13** | BCN-391 | Spain | M | 75 | A2B1C1 | - | - | - | - | - |
| **MSA 14** | BCN-427 | Spain | F | 81 | A2B1C2 | - | - | - | - | - |
| **MSA 15** | BCN-480 | Spain | M | 70 | A0B1C0 | - | - | - | - | - |

| Case | Code | Source | Sex | Age | NIA-AA | Cerebral Amyloid Angiopathy | Lewy Body Pathology | Lewy Body Braak Stage | LATE-NC Stage | Other Pathologic Diagnosis |
|---|---|---|---|---|---|---|---|---|---|---|
| **PSP 1** | 1014 | Canada | F | 77 | - | Aβ-positive Type 2 | - | Amygdala only | - | - |
| **PSP 2** | 1443 | Canada | F | 74 | - | - | - | - | - | - |
| **PSP 3** | 1552 | Canada | F | 74 | - | - | - | - | - | - |
| **PSP 4** | R-NBC-19-1 | Canada | F | 70 | - | - | - | - | - | - |
| **PSP 5** | R-NBC-20-13 | Canada | M | 73 | - | Aβ-positive Type 2 | - | - | - | Argyrophilic Grain Disease (4R) |
| **Control 1** | 1051 | Canada | M | 74 | - | - | - | - | - | - |
| **Control 2** | 1095 | Canada | F | 48 | - | - | - | - | - | - |
| **Control 3** | 1113 | Canada | F | 69 | - | - | - | - | - | - |
| **Control 4** | 1466 | Canada | F | 53 | - | - | - | - | - | - |
| **Control 5** | 1509 | Canada | F | 26 | - | - | - | - | - | - |
